# Emotion Schemas are Embedded in the Human Visual System

**DOI:** 10.1101/470237

**Authors:** Philip A. Kragel, Marianne Reddan, Kevin S. LaBar, Tor D. Wager

**Affiliations:** Department of Psychology and Neuroscience and the Institute of Cognitive Science, University of Colorado, Boulder, CO, USA; Institute for Behavioral Genetics, the University of Colorado, Boulder, CO, USA; Department of Psychology and Neuroscience and the Center for Cognitive Neuroscience, Duke University, Durham, NC, USA

## Abstract

Theorists have suggested that emotions are canonical responses to situations ancestrally linked to survival. If so, then emotions may be afforded by features of the sensory environment. However, few computationally explicit models describe how combinations of stimulus features evoke different emotions. Here we develop a convolutional neural network that accurately decodes images into 11 distinct emotion categories. We validate the model using over 25,000 images and movies and show that image content is sufficient to predict the category and valence of human emotion ratings. In two fMRI studies, we demonstrate that patterns of human visual cortex activity encode emotion category-related model output and can decode multiple categories of emotional experience. These results suggest that rich, category-specific emotion representations are embedded within the human visual system.

## Introduction

Emotions are thought to be canonical responses to situations ancestrally linked to survival (Tooby and Cosmides) or the well-being of an organism (Lazarus 1968). Sensory processing plays a prominent role in nearly every theoretical explanation of emotion (e.g., Scherer 1984, Ekman 1992, Russell 2003, Niedenthal 2007, Barrett 2017), yet neuroscientific views have historically suggested that emotion is driven by specialized brain regions, e.g., in the limbic system (MacLean 1952) and related subcortical circuits (Panksepp 1998), or in some theories in neural circuits specialized for emotion categories such as fear (Adolphs 2013) and sadness (Mayberg, Liotti et al. 1999). According to these longstanding views, activity in sensory cortex (e.g., visual areas V1-V4) is thought to be *antecedent* to emotion, but not central to emotional appraisals, feelings, or responses. However, recent theoretical developments (Pessoa 2008, Barrett and Bar 2009, Pessoa and Adolphs 2010) and empirical observations suggest that sensory and emotional representations may be much more intertwined than previously thought. Activity in visual cortex is enhanced by emotionally significant stimuli (Morris, Friston et al. 1998, Vuilleumier, Richardson et al. 2004), and single neurons learn to represent the affective significance of stimuli. For example, neurons in V1 (Shuler and Bear 2006), V4 (Haenny and Schiller 1988), and inferotemporal cortex (Mogami and Tanaka 2006, Eradath, Mogami et al. 2015, Sasikumar, Emeric et al. 2018) selectively respond to rewarding stimuli. In addition, multivariate patterns of human brain activity that predict emotion-related outcomes often utilize information encoded in visual cortex (Chang, Gianaros et al. 2015, Kragel and LaBar 2015, Krishnan, Woo et al. 2016, Saarimaki, Gotsopoulos et al. 2016, Saarimaki, Ejtehadian et al. 2018).

There are at least two ways of interpreting this evidence. On one hand, emotion-related activity in sensory areas could reflect a general enhancement of visual processing for relevant, novel, or attended percepts (O’Connor, Fukui et al. 2002, McAlonan, Cavanaugh et al. 2008). Stronger sensory responses to emotionally relevant percepts can also be evolutionarily conserved (relevant in ancestral environments (Öhman and Mineka 2003)) or learned during development (Held and Hein 1963, Recanzone, Schreiner et al. 1993, Shuler and Bear 2006). In this case, affective stimuli evoke stronger sensory responses, but the information about emotion content (fear vs. anger, sadness vs. joy) is thought to be represented elsewhere. Alternatively, perceptual representations in sensory (e.g., visual) cortex could reflect the *content* of emotional responses in a rich way; specific configurations of perceptual features could afford specific types, or categories, of emotional responses, including fear, anger, desire, joy, etc. In this case, neural codes in sensory cortices might represent information directly relevant for the nature of emotional feelings and responses.

The latter view is broadly compatible with appraisal theories (Moors 2018) and more recent theories of emotions as constructed from multiple perceptual, mnemonic, and conceptual ingredients (Russell 2003, Barrett 2006, Barrett 2017). In the former, *emotion schemas* (Izard 2007) are canonical patterns of organism-environment interactions that afford particular emotions. For example, scenes of carnage evoke rapid responses related to disgust or horror, and later (integrating conceptual beliefs about the actors and other elements), compassion, anger, or other emotions. Scenes with attractive, scantily clad people evoke schemas related to sex; scenes with delicious food evoke schemas related to consumption; and so on. In these cases, the sensory elements of the scene do not fully determine the emotional response—other ingredients are involved, including one’s personal life experiences, goals and interoceptive states (Bower 1981, Izard 2007)—but the sensory elements *are* sufficient to convey the schema or situation that the organism must respond to. Initial appraisals of emotion schemas (often called “System 1” appraisals) can be made rapidly (Lazarus 1966, Kahneman and Egan 2011) and in some cases unconsciously, and unconscious emotion may drive preferences and shape learning (Zajonc 1984, Berridge and Winkielman 2003, Pessiglione, Seymour et al. 2006). Emotion schemas are also content-rich in the sense that they sharply constrain the repertoire of emotional responses afforded by a given schema. For example, horror scenes might afford fear, anger, or compassion, but other kinds of emotional responses (sadness, nurturing, playfulness) would be ancestrally inappropriate. Thus, while some affective primitives (representations related to survival and well-being) are related to biologically older subcortical brain systems (MacLean 1952, Panksepp 1998) and involve relatively little cognitive processing (Ekman and Cordaro 2011), canonical, category-specific *emotion schemas* exist and may be embedded in part in human sensory cortical systems.

The hypothesis that emotion schemas are embedded in sensory systems makes several predictions that have not, to our knowledge, been tested. First, models constructed from image features *alone* should be able to (a) predict normative ratings of emotion category made by humans and (b) differentiate *multiple emotion categories*. Second, representations in such models should map onto distinct patterns of brain activity in sensory (i.e., visual) cortices. Third, sensory areas, and particularly visual cortex, should be sufficient to decode multiple emotion categories. Here, we test each of these hypotheses.

To test Predictions 1 and 2, we developed a convolutional neural network (CNN) whose output is a probabilistic representation of the emotion category of a picture or video, and used it to classify images into 20 different emotion categories using a large stimulus set of 2,232 emotional video clips (Cowen and Keltner 2017). We validated this model, called EmoNet, in three different contexts, by predicting: (i) normative emotion categories of video clips not used for training; (ii) normative emotional intensity ratings for International Affective Picture System (IAPS), an established set of emotional images (Lang, Bradley et al. 2008); and (iii) the genre of cinematic movie trailers, which are designed to manipulate emotion by presenting different visual cues (Rasheed and Shah 2002). To test whether EmoNet can uniquely identify multiple emotion categories, we developed and applied a statistical framework for estimating the number of discriminable emotion categories. To test Prediction 2, we used machine learning approaches to find patterns of brain activity in the occipital lobe (measured via fMRI, *N* = 18) linked to emotion category-related output from EmoNet. To test Prediction 3, in a separate fMRI study (*N* = 32), we verified that patterns of occipital lobe activity can decode the category of emotional responses elicited by videos and music (across 5 categories). Our results are consistent with prior research showing that different patterns of visual cortical activity are associated with different emotion categories (Chang, Gianaros et al. 2015, Kragel and LaBar 2015, Krishnan, Woo et al. 2016, Saarimaki, Gotsopoulos et al. 2016, Saarimaki, Ejtehadian et al. 2018), but goes beyond them to (1) rigorously test whether sensory representations are sufficient for accurate decoding, and (2) provide a computationally explicit account of how sensory inputs are transformed into emotion-related codes.

## Classifying visual images into multiple emotion categories

EmoNet (**Figure 1**) was based on the popular AlexNet image recognition model, and used representations learned from AlexNet as input into a final fully-connected layer trained to predict the normative emotion category of over 137,482 images extracted from videos (Cowen and Keltner 2017) with normative emotion categories were based on ratings from 853 subjects. We tested EmoNet on 24,634 images from 400 videos not included in the training set. EmoNet accurately decoded normative human ratings of emotion catgories, providing support for Prediction 1. The human-consensus category was among the top 5 predictions made by the model (top-5 accuracy in 20-way classification) for 62.6% of images (chance = 27.95%; *P* < .0001, permutation test); the top-1 accuracy in a 20-way classification was 23.09% (chance = 5.00%; *P* < .0001, permutation test); the average area under the receiver operating characteristic curve across the 20 categories was .745 (Cohen’s *d* = 0.945), indicating that emotions could be discriminated from one another with large effect sizes.

**Figure 1.**
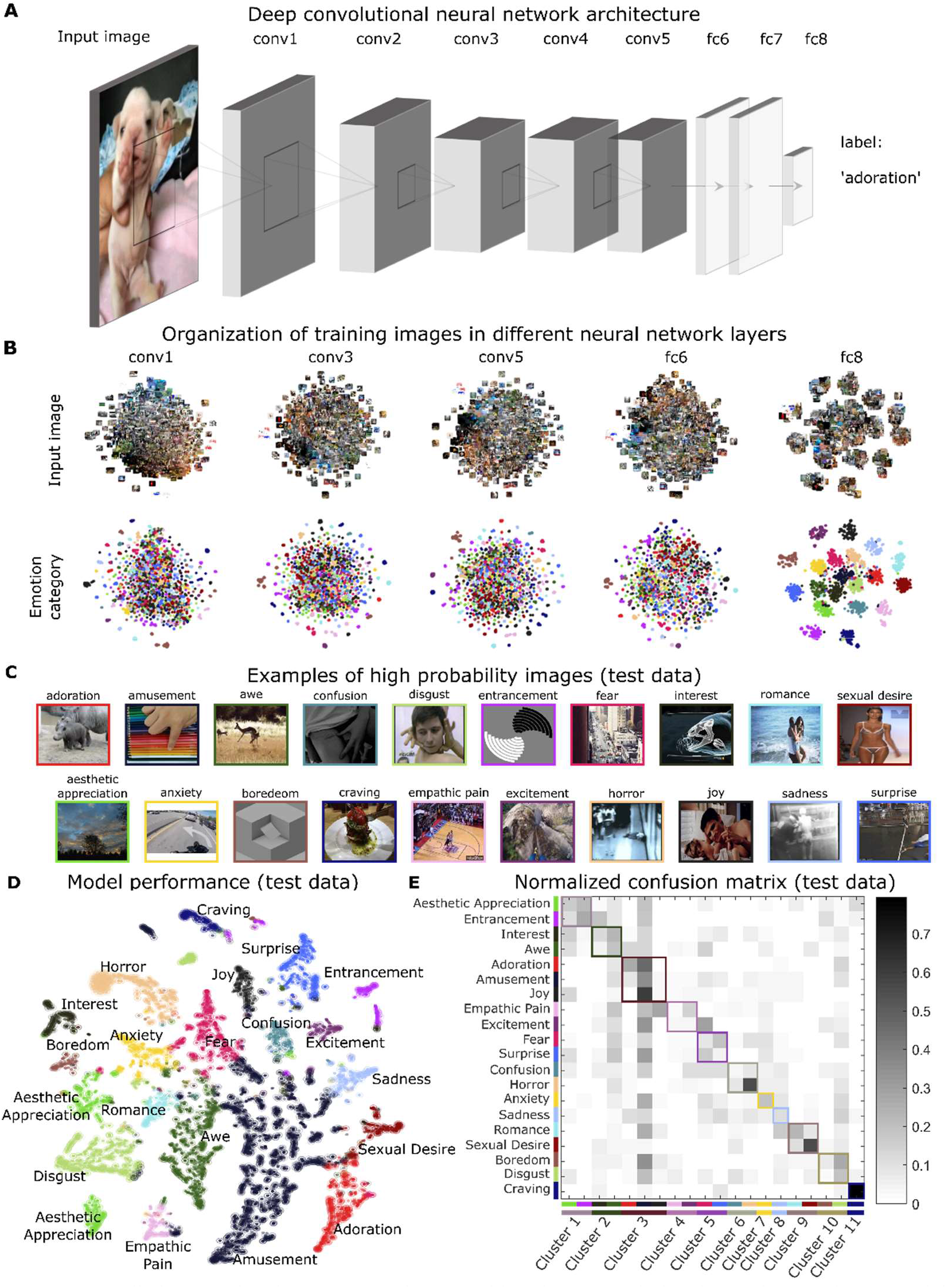
Predicting emotional responses to images with a deep convolutional neural network. (a) Model architecture follows that of AlexNet (five convolutional layers followed by three fully-connected layers), only the last fully-connected layer has been retrained to predict emotion categories. (b) Activation of artificial neurons in three convolutional layers (1, 3, and 5) and the last two fully-connected layers (6 and 8) of the network. Scatterplots depict t-distributed Stochastic Neighbor Embedding (*t*-SNE) plots of activation for a random selection of 1,000 units in each layer. The first four layers come from a model developed to perform object-recognition (Krizhevsky, Sutskever et al. 2012), and the last-layer was retrained to predict emotion categories from an extensive database of video clips. Note the progression away from low-level image features towards more abstract emotion schemas. (c) Examples of randomly selected images assigned to each class in hold-out test data (images from videos that were not used for training the model). Pictures were not chosen to match target classes. Some examples show contextually-driven prediction, e.g., an image of a sporting event is classified as ‘empathic pain’ even though no physical injury is apparent. (d) t-SNE plot shows model predictions in test data. Colors indicate the predicted class, and circled points indicate that the ground truth label was in the top five predicted categories. Although t-SNE does not preserve global distances, the plot does convey local clustering of emotions such as ‘amusement’ and ‘adoration.’ (e) Normalized confusion matrix shows the proportion of test data that are classified into the twenty categories. Rows correspond to the correct category of test data, and columns correspond to predicted categories. Gray colormap indicates the proportion of predictions in the test dataset, where each row sums to a value of 1. Correct predictions fall on the diagonal of the matrix, whereas erroneous predictions comprise off-diagonal elements. Categories the model is biased towards predicting, such as ‘amusement,’ are indicated by dark columns. Data-driven clustering of errors shows 11 groupings of emotions that are all distinguishable from one another (see Materials and Methods and Figure S1).

Crucially, EmoNet accurately discriminated *multiple emotion categories* in a relatively fine-grained way, though model performance varied across categories. ‘Craving’ (AUC = .987, 95% CI = [.980 .990]; *d* = 3.13; *P* < .0001), ‘sexual desire’ (AUC = .965, 95% CI = [.960 .968]; *d* = 2.56; *P* < .0001), ‘entrancement’ (AUC = .902, 95% CI = [.884 .909]; *d* = 1.83; *P* < .0001), and ‘horror’ (AUC = .876, 95% CI = [.872 .883]; *d* = 1.63; *P* < .0001) were the most accurately predicted categories. On the other end of the performance spectrum, ‘confusion’ (AUC = .636, 95% CI = [.621 .641]; *d* = .490; *P* < .0001), ‘awe’ (AUC = .615, 95% CI = [.592 .629]; *d* = .415; *P* < .0001), and ‘surprise’ (AUC = .541, 95% CI = [.531 .560]; *d* = .147; *P* = .0002) exhibited the lowest levels of performance, despite exceeding chance levels. Some emotions were highly confusable in the test data, such as ‘amusement’, ‘adoration’, and ‘joy’, suggesting they have similar visual features despite being distinct from other emotions (**Figure S1**). Thus, visual information is sufficient for predicting some emotion schemas, particularly those that have a strong relationship with certain high-level visual categories, such as ‘craving’ or ‘sexual desire’, whereas other sources of information are necessary to discriminate emotions that are conceptually abstract or depend on temporal dynamics (e.g., ‘confusion’ or ‘surprise’).

To further assess the number of distinct emotion categories represented by EmoNet, we developed two additional tests of (1) dimensionality and (2) emotion category discriminability. First, we tested the possibility that EmoNet is tracking a lower-dimensional space, such as one organized by *valence* and *arousal* (Russell 1980), rather than a rich category-specific representation. Principal components analysis (PCA) on model predictions in the hold-out dataset indicated that many components were required to explain model predictions; 17 components were required to explain 95% of the model variance, with most components being mapped to only a single emotion (i.e., exhibiting simple structure (Carroll 1953), see **Figure S1**). To test category discriminability, we developed a test of how many emotion categories were *uniquely discriminable from each other category* in EmoNet’s output (**Figure 1E;** see Supplementary Text for details of the method). The results indicated that EmoNet differentiated 11 (95% CI = [10 to 14]) distinct emotion categories from one another, supporting the sensory embedding hypothesis.

## Modeling valence and arousal as combinations of emotion-related features

To further test EmoNet’s generalizability, we tested it on three additional image and movie databases. A first test applied EmoNet to images in the International Affective Picture System (IAPS), a widely studied set of images used to examine the influence of positive and negative affect on behavior, cognitive performance, autonomic responses, and brain activity (Lang and Bradley 2007). The IAPS dataset provides an interesting test because human norms for emotion intensity ratings are available, and because IAPS images often elicit mixed emotions that include responses in multiple categories (Mikels, Fredrickson et al. 2005). Much of the variance in these emotion ratings is explained by a two-dimensional model of valence (pleasant to unpleasant) and arousal (calm to activated), and emotion categories are reliably mapped into different portions of the valence-arousal space (Bradley and Lang 1999, Tellegen, Watson et al. 1999, Fontaine, Scherer et al. 2007, Warriner, Kuperman et al. 2013), often in a circumplex pattern (Russell 1980, Plutchik 1997, Russell and Barrett 1999, Cowen and Keltner 2017). These features allowed us to assess whether EmoNet predicts normative human ratings of valence and arousal across the full IAPS dataset, and whether EmotNet organizes emotions in a low-dimensional or circumplex structure similar to human ratings.

We constructed predictive models using partial least squares (PLS) regression of human valence and arousal on features from the last fully connected layer of EmoNet, which has 20 units, one for each emotion category. We analyzed the accuracy in predicting valence and arousal ratings of out-of-sample test images using 10-fold cross-validation (Kohavi 1995), stratifying folds based on normative ratings. We also analyzed the model weights (*β_PLS_*) mapping emotion categories to arousal and valence, to construct a valence and arousal space from the activity of emotion category units in EmoNet. The models strongly predicted valence and arousal ratings for new (out-of-sample) images. The model predicted valence ratings with *r* = .88 (*P* < .0001, permutation test, RMSE = 0.9849), and arousal ratings with *r* = .85 (*P* < .0001, RMSE = 0.5843). A follow-up generalization test using these models to predict normative ratings on a second, independent image database (Kurdi, Lozano et al.2017)—with no model retraining—showed similar levels of performance for both valence (*r* = .83, *RMSE* = 1.605) and arousal (*r* = .84, *RMSE* = 1.696). Thus, EmoNet explained over 60% of the variance in average human ratings of pleasantness and arousal when viewing IAPS images. This level of prediction indicate that EmoNet predicts valence and arousal in stimuli that elicit mixed emotions.

**Figure 2.**
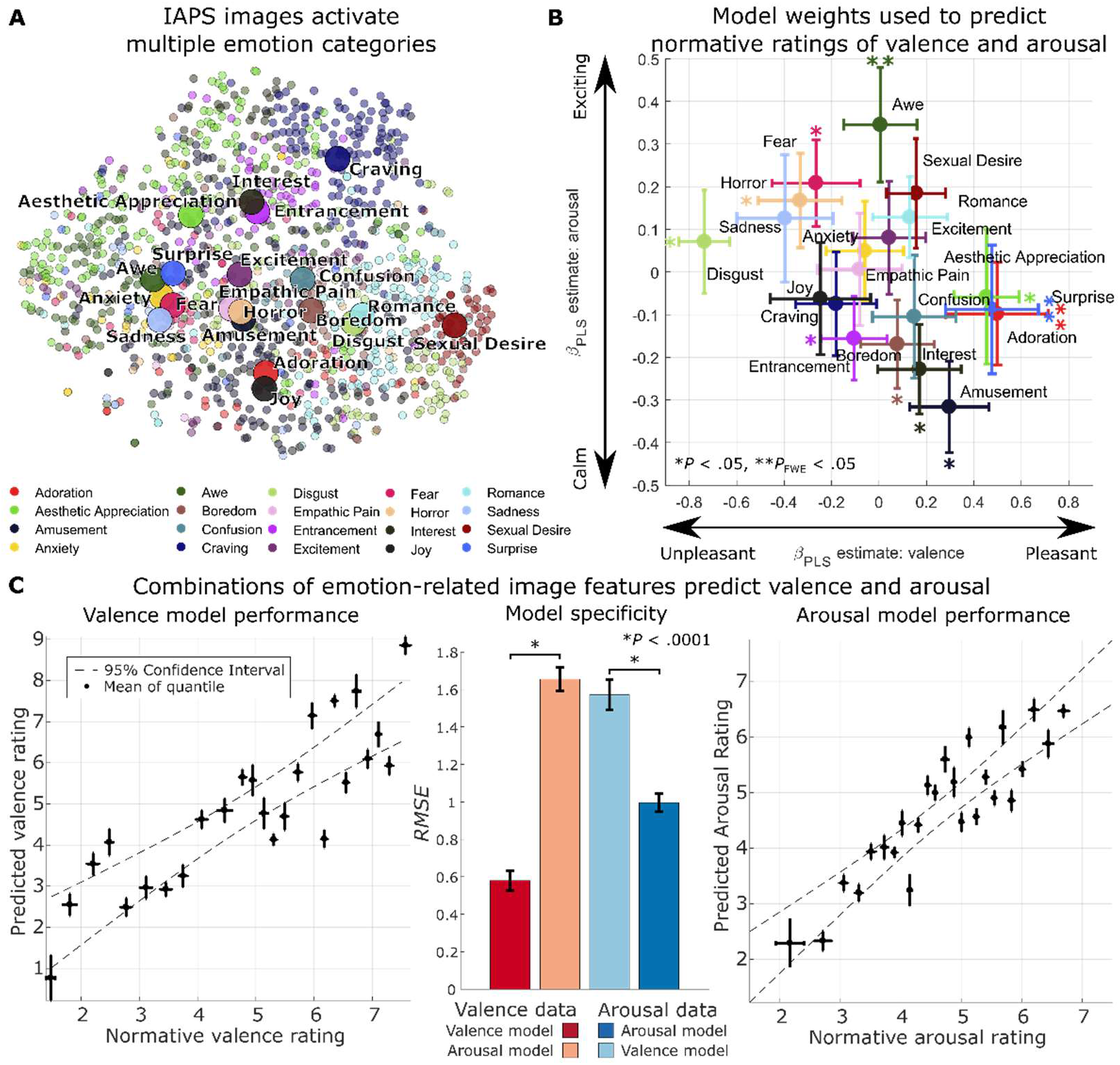
Emotion-related image features predict normative ratings of valence and arousal. (a) Depiction of the full International Affective Picture System (IAPS), with picture locations determined by *t*-Distributed Stochastic Neighbor Embedding of activation of the last fully connected layer of EmoNet. The color of each point indicates the emotion category with the greatest score for each image. Large circles indicate mean location for each category. Combinations of loadings on different emotion categories are used to make predictions about normative ratings of valence and arousal. (b) Parameter estimates indicate relationships identified using Partial least squares regression to link the 20 emotion categories to the dimensions of valence (x-axis) and arousal (y-axis). Bootstrap means and standard errors are shown by circles and error bars. For predictions of valence, positive parameter estimates indicate increasing pleasantness, and negative parameter estimates indicate increasing unpleasantness; for predictions of arousal, positive parameter estimates indicate a relationship with increasing arousal and negative estimates indicate a relationship with decreasing arousal. \**P* < .05, \*\**P*_FWE_ < .05 (c) Cross-validated model performance. Left and right panels show normative ratings of valence and arousal, plotted against model predictions. Individual points reflect the average rating for each of 25 quintiles of the full IAPS set. Error bars indicate the standard deviation of normative ratings (x-axis, *N* = 47) and the standard deviation of repeated 10-fold cross-validation estimates (y-axis, *N* = 10). Bar plots in the middle panel show overall root-mean-square error (*RMSE*, lower values indicate better performance) for models tested on valence data (left bars, red hues) and arousal data (right bars, blue hues). Error bars indicate the standard deviation of repeated 10-fold cross-validation. \**P* < .0001 corrected resampled *t*-test. The full convolutional neural network model and weights for predicting valence and arousal are available at https://github.com/canlab for public use.

In addition, the categorical emotion responses in EmoNet’s representation of each image were arranged in valence-arousal space in a manner similar to the human circumplex model (Russell 1980) (**Figure 2b**), though with some differences from human raters. Units coding for ‘adoration’ (*β̂*= .5002, 95% CI = [.2722 1.0982]), ‘aesthetic appreciation’ (*β̂*= .4508, 95% CI = [.1174 .6747]), and ‘surprise’ (*β̂*= .4781, 95% CI = [.3027 1.1476]) were most strongly associated with positive valence across categories. Units coding for ‘disgust’ (*β̂*= −.7377, 95% CI = [−1.0365 −.6119]), ‘entrancement’ (*β̂*= −.1048, 95% CI = [−.5883 −.0010]), and ‘horror’ (*β̂*= −.3311, 95% CI = [−.7591 −.0584]) were the most negatively valenced. The highest loadings on arousal were in units coding for ‘awe’ (*β̂*= .0285, 95% CI = [.0009 .0511]) and ‘horror’ (*β̂*= .0322, 95% CI = [.0088 .0543]), and the lowest-arousal categories were ‘amusement’ (*β̂*= −.3189, 95% CI = [−.5567 −.1308]), ‘interest’ (*β̂*= −.2310, 95% CI = [−.4499 −.0385]) and ‘boredom’ (*β̂*= −.1605, 95% CI = [−.4380 −.0423]). The marked similarities with the human affective circumplex demonstrate that model representations of emotion categories reliably map onto dimensions of valence and arousal. However, these findings do *not* indicate that the valence-arousal space is sufficient to encode the full model output; in fact, we estimate that doing so requires 17 dimensions, and the loadings in Figure 2b do not exhibit a classic circumplex pattern. The discrepancies (e.g., ‘surprise’ is generally considered high-arousal and neutral valence, and ‘awe’ is typically positively valenced) highlight that the model was designed to track visual features that might serve as ingredients of emotion, but do not capture human feelings in all respects. For example, while people may typically rate ‘awe’ as a positive experience, ‘awe-inspiring’ scenes often depict high-arousal activities (e.g., extreme skiing or base jumping).

## Classifying the genre of movie trailers based on their emotional content

A second test examined whether emotion categories could meaningfully be applied to dynamic stimuli such as videos. We tested EmoNet’s performance in classifying the genre of 28 randomly sampled movie trailers from romantic comedy (*N* = 9), action (*N* = 9), and horror (*N* = 10) genres (see Materials and Methods for sampling and selection criteria). EmoNet made emotion predictions for each movie frame (for an example see the time series in **Figure 3**). PLS regression was used to predict movie genres from the average activation over time in EmoNet’s final emotion category layer, using one-vs-all classification (Rifkin and Klautau 2004) with 10-fold cross-validation to estimate classification accuracy in independent movie trailers.

The results indicated that EmoNet’s frame-by-frame predictions tracked meaningful variation in emotional scenes across time (**Figure 3a**), and that mean emotion category probabilities accurately classified the trailers (**Figure 3b-c**), with three-way classification accuracy of 71.43% (*P* < .0001, permutation test; chance is 35.7%). The average area under the receiver operating characteristic curve for the three genres was .855 (Cohen’s *d* = 1.497; **Figure 3c**). Classification errors were made predominantly between action and horror movies (26.32%), whereas romantic comedies were not misclassified, indicating that they had the most distinct features.

**Figure 3.**
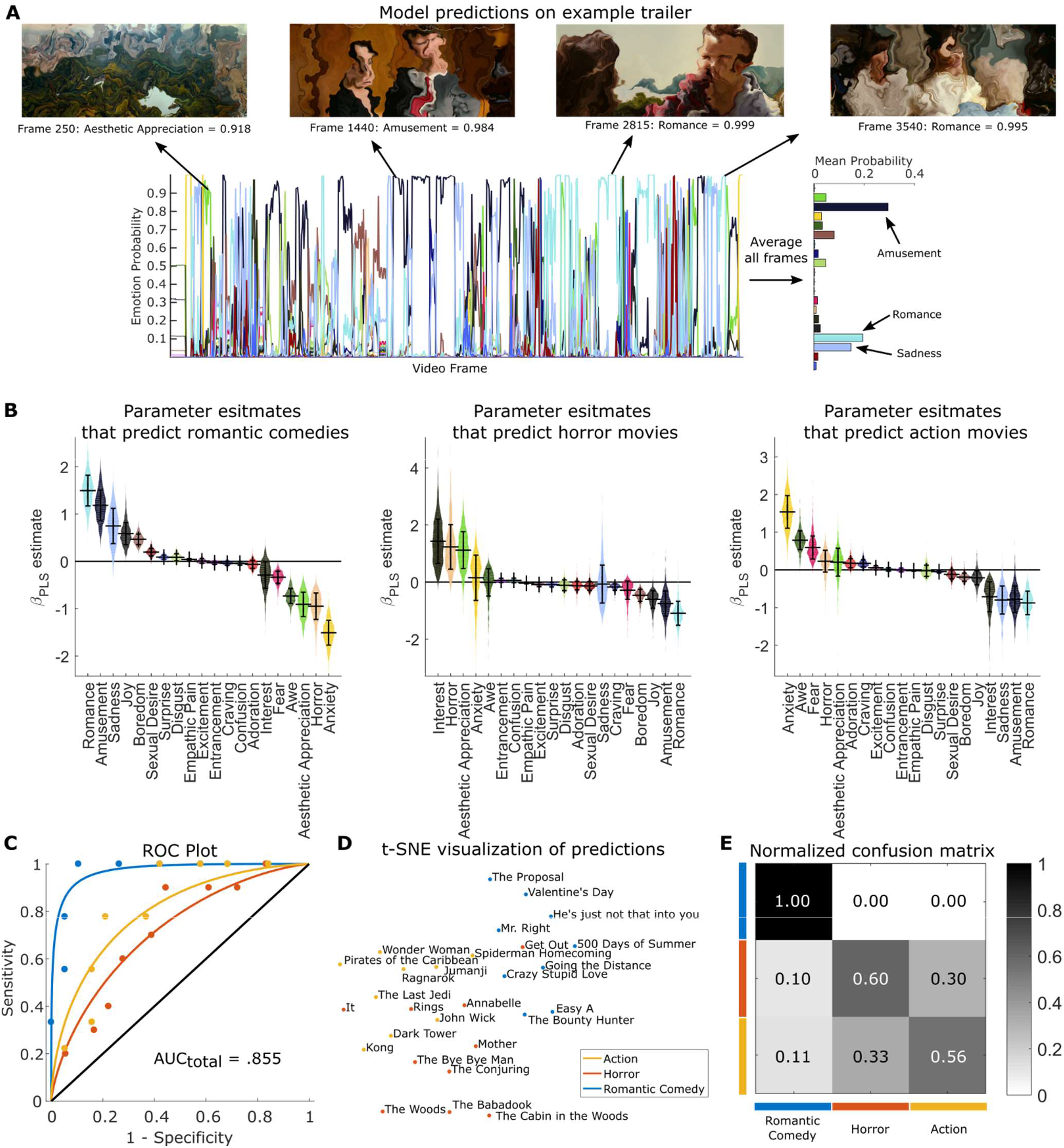
Identifying the genre of movie trailers using emotional image features. (a) Emotion prediction for a single movie trailer. Time-courses indicate model outputs on every fifth frame of the trailer for the twenty emotion categories, with example frames shown above. A summary of the emotional content of the trailer is shown on the right, which is computed by averaging predictions across all analyzed frames. (b) Partial least squares parameter estimates indicate which emotions lead to predictions of different movie genres. Violin plots depict the bootstrap distributions (1,000 iterations) for parameters estimates differentiating each genre from all others. Error bars indicate bootstrap standard error. (c) Receiver operator characteristic plots depict 10-fold cross-validation performance for classification. The solid black line indicates chance performance. (d) *t*-SNE plot based on the average activation of all 20 emotions. (e) Confusion matrix depicting misclassification of different genres; rows indicate the ground truth label and columns indicate predictions. The grayscale color bar shows the proportion of trailers assigned to each class.

Movie genres are systematically associated with different emotion schemas: romantic comedies were predicted by increased activation of units coding for ‘romance’ (*β̂*= 1.499, 95% CI = [1.001 2.257]), ‘amusement’ (*β̂*= 1.167, 95% CI = [0.639 2.004]), and ‘sadness’ (*β̂*= 0.743, 95% CI = [0.062 1.482]); horror trailers were predicted by activation of ‘interest’ (*β̂*= 1.389, 95% CI = [0.305 3.413]), ‘horror’ (*β̂*= 1.206, 95% CI = [0.301 3.536]), and ‘aesthetic appreciation’ (*β̂*= 1.117, 95% CI = [0.259 2.814]); and action trailers were predicted by activation of ‘anxiety’ (*β̂*= 1.526, 95% CI = [0.529 2.341]), ‘awe’ (*β̂*= 0.769, 95% CI = [0.299 1.162]), and ‘fear’ (*β̂*= 0.575, 95% CI = [0.094 1.109]). As with IAPS images, EmoNet tracked canonical visual scenes that can lead to several kinds of emotional experience based on context. For instance, some horror movies in this sample included scenic shots of woodlands, which were classified as ‘aesthetic appreciation’, leading to high weights for ‘aesthetic appreciation’ on horror films. While such mappings illustrate how EmoNet output alone should not be over-interpreted in terms of human feelings, they also illustrate how emotion concepts can constrain the repertoire of feelings-in-context. A beautiful forest or children playing can be ominous when paired with other threatening context cues (e.g., scary music), but the emotion schema is incompatible with a range of other emotions (sadness, anger, interest, sexual desire, disgust, etc.).

## Decoding model representations of emotions from patterns of human brain activity

If emotion schemas are afforded by visual scenes, then it should be possible to decode emotion category-related representations in EmoNet from activity in the human visual system. To test this hypothesis, we measured brain activity using fMRI while participants (*N* = 18) viewed a series of 112 affective images that varied in affective content (see Materials and Methods for details). Treating EmoNet as a model of the brain (Yamins and DiCarlo 2016), we used PLS to regress patterns in EmoNet’s emotion category layer onto patterns of fMRI responses to the same images (e.g., see (Yamins, Hong et al. 2014) for an application of this approach to object recognition). We investigated the predictive performance, discriminability, and spatial localization of these mappings to shed light on how and where emotion-related visual scenes are encoded in the brain.

Because EmoNet was trained on visual images, we first explored how emotion schemas might emerge from activity in the human visual system, within a mask comprising the entire occipital lobe (7,214 voxels (Lancaster, Woldorff et al. 2000)). Patterns of occipital activity predicted variation in EmoNet’s emotion category units across images, with different fMRI patterns associated with different emotion categories (**Figure 4a**, for individual maps, see **Figure S2**). Multiple correlations between brain-based predictions and activation in EmoNet emotion category units were tested in out-of-sample individuals using leave-one-subject-out (Esterman, Tamber-Rosenau et al. 2010) cross-validation. These correlations were positive and significant for each of the 20 EmoNet emotion categories (mean *r* = 0.2819 ± .0163 (*SE*) across subjects, mean effect size *d* = 3.00, 76.93% of the noise ceiling, *P* < .0001, permutation test, see Supplementary Text). The highest average level of performance included ‘entrancement’ (*r* = 0.4537 ± .0300 (*SE*), *d* = 3.559, 77.03% of the noise ceiling, *P* < .0001), ‘sexual desire’ (*r* = 0.4508 ± .0308 (*SE*), *d* = 3.453, 79.01% of the noise ceiling, *P* < .0001), and ‘romance’ (*r* = 0.3861 ± .0203 (*SE*), *d* = 4.476, 72.34% of the noise ceiling, *P* < .0001), whereas ‘horror’ (*r* = 0.1890 ± .0127 (*SE*), *d* = 3.520, 60.17% of the noise ceiling, *P* < .0001), ‘fear’ (*r* = 0.1800 ± .0216 (*SE*), *d* = 1.963, 59.44% of the noise ceiling, *P* < .0001), and ‘excitement’ (*r* = 0.1637 ± .0128 (*SE*), *d* = 3.004, 65.28% of the noise ceiling, *P* < .0001), exhibited the lowest levels of performance.

**Figure 4.**
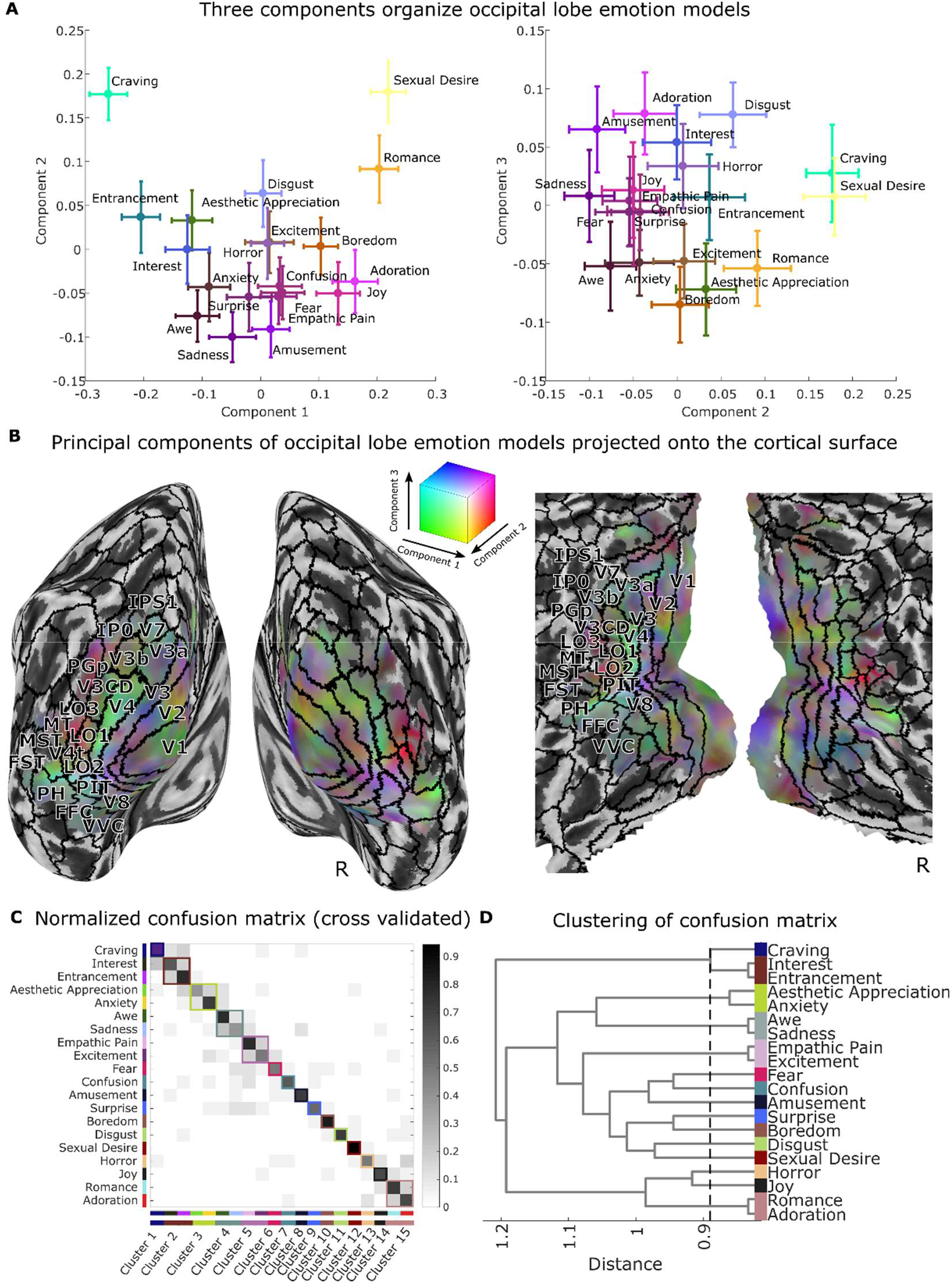
Visualization of the 20 occipital lobe models using principal components analysis (PCA) reveals three important emotion-related features of the visual system. (a) Scatter plots depict the location of 20 emotion categories in PCA space, with colors indicating loadings onto the first three principal components (PC_s_) identified from 7,214 voxels that retain approximately 95% of the spatial variance across categories. The color of each point is based on the component scores for each emotion (in an additive red-green-blue color space, PC_1_ = red, PC_2_ = green, PC_3_ = blue). Error bars reflect bootstrap standard error. (b) Visualization of group average coefficients that show mappings between voxels and principal components. Colors are from the same space as depicted in panel (a). Solid black lines indicate boundaries of cortical regions based on a multi-modal parcellation of the cortex (Glasser, Coalson et al. 2016). Surface mapping and rendering were performed using the CAT12 toolbox (Dahnke, Yotter et al. 2013, Gaser and Dahnke 2016). (c) Normalized confusion matrix shows the proportion of data that are classified into 20 emotion categories. Rows correspond to the correct category of cross validated data, and columns correspond to predicted categories. Gray colormap indicates the proportion of predictions in the dataset, where each row sums to a value of 1. Correct predictions fall on the diagonal of the matrix; erroneous predictions comprise off-diagonal elements. Data-driven clustering of errors shows 15 groupings of emotions that are all distinguishable from one another. (d) Visualization of distances between emotion groupings. Dashed line indicates minimum cutoff that produces 15 discriminable categories. Dendrogram was produced using Ward’s linkage on distances based on the number of confusions displayed in panel (c). See Supplementary Text for a description and validation of the method.

To further test the number of discriminable emotion categories encoded in visual cortex, we constructed a confusion matrix for relationships between the visual cortical multivariate pattern responses and EmoNet emotion category units. For each study participant, we correlated the output from each of the 20 fMRI models (a vector with 112 values, one for each IAPS image) with vectors of activation across EmoNet’s 20 emotion category units (producing a 20 × 20 correlation matrix), using leave-one-subject out cross validation to provide an unbiased test. For each model, the EmoNet unit with the highest correlation was taken as the best-guess emotion category based on brain activity, and the resulting confusion matrix was averaged across participants. The confusion matrix is shown in **Figure 4c**, with correct predictions in 20-way classification (sensitivity) shown on the diagonal, and false alarms (1 – specificity) on the off-diagonal. The average sensitivity across subjects was 66.67 ± 11.4% (*SEM*) and specificity was 97.37 ± .88%; thus, visual cortical activity was mapped onto EmoNet’s categories with a positive predictive value of 65.45 ± 10.4% (chance is approximately 5%). In addition, as above, we estimated the number of uniquely discriminable categories by clustering the 20 the categories and searches the clustering dendrogram to determining the maximum number of clusters (minimum link distance) at which each cluster was significantly discriminable from each other one, with bootstrap resampling to estimate confidence intervals. The results showed at least 15 discriminable categories (95% CI = [15 17]), with a pattern of confusions that was sometimes intuitive based on psychology (e.g., empathic pain was indistinguishable from excitement, romance was grouped with adoration and interest with entrancement), but in other cases was counterintuitive (sadness grouped with awe). This underscores that the visual cortex does not perfectly reproduce human emotional experience, but nonetheless contain a rich, multidimensional representation of high-level, emotion-related features, in support of Prediction 2.

In additional model comparisons, we tested whether occipital cortex was necessary and sufficient for accurate prediction of EmoNet’s emotion category representation. We compared models trained using brain activity from individual areas (i.e., V1-V4 (Amunts, Malikovic et al. 2000, Rottschy, Eickhoff et al. 2007) and inferotemporal cortex (Tzourio-Mazoyer, Landeau et al. 2002)), the entire occipital lobe (Lancaster, Woldorff et al. 2000), and the whole brain. We trained models to predict variation across images in each EmoNet emotion category unit, and averaged performance across emotion categories. The whole-occipital-lobe model (*r* = 0.2819 ± .0163 (*SE*)) and the whole-brain model (*r* = 0.2664 ± .0150 (*SE*)) predicted EmoNet emotion categories more strongly than models based on individual visual areas (*r* = .0703 to 0.1635, all *P* < .0001). Furthermore, the occipital lobe model showed marginally better performance than the whole-brain model (Δ*r* = .0155, *P* = .0404, 95% CI = [0.0008 0.0303], paired t-test), despite having nearly 100,000 fewer features available for prediction (**Figure S3**). A post hoc, confirmatory test revealed that excluding occipital lobe activation from the whole-brain model significantly reduced performance (Δ*r* = −.0240, *P* < .0001, 95% CI = [−0.0328 −0.0152], paired t-test), indicating that activity in the occipital lobe meaningfully contributed to predictions in the whole-brain model. These results provide strong support for distributed representation of emotion schemas within the occipital lobe and partially redundant coding of this information in other brain systems. The distributed codes for emotion categories parallel other recent findings on population coding of affective processes (Chang, Gianaros et al. 2015, Krishnan, Woo et al. 2016); for review, see ref. (Kragel, Koban et al. 2018).

## Classifying patterns of visual cortex activity into multiple distinct emotion categories

To provide additional evidence that visual cortical representations are emotion category-specific, we tested whether visual cortical activity was sufficient to decode the category of emotional videos in an independent dataset (N = 32; see ref. Kragel and LaBar 2015). In this dataset, human subjects viewed cinematic film clips that elicited contentment, sadness, amusement, surprise, fear, and anger. Videos were selected that elicited responses in one emotion category above all others for each video, complementing the previous study, whose stimuli elicited more blended emotional responses. We tested predictive accuracy in seven-way classification of emotion category based on subject-average patterns of occipital lobe activity for each condition, with eight-fold cross-validation across participants to test prediction performance in out-of-sample individuals. We then performed discriminable cluster identification (**Figures 1** and **4**, see Supplementary Text for details) to estimate how many distinct emotion categories out of this set are represented in visual cortex.

This analysis revealed that of the seven states being classified (six emotions and neutral videos), at least five distinct emotion clusters (95% CI = [5 7]) could be reliably discriminated from one another based on occipital lobe activity (5-way classification accuracy = 40.54%, chance = 20% see **Figure 5**), supporting Prediction 3. Full seven-way classification was 29.95% (chance = 14.3%, *P* = 0.002). Contentment, amusement, and neutral videos were reliably differentiated from all other emotions. States of fear and surprise were not discriminable from one another (they were confused 21.09% of the time), yet they were reliably differentiated from all other emotions. Sadness and anger were also confusable (15.5%) but were discriminable from all other emotional states. Thus, although some emotional states were similar to one another regarding occipital lobe activation, we found strong evidence for categorical coding of multiple emotions during movie inductions of specific emotions.

**Figure 5.**
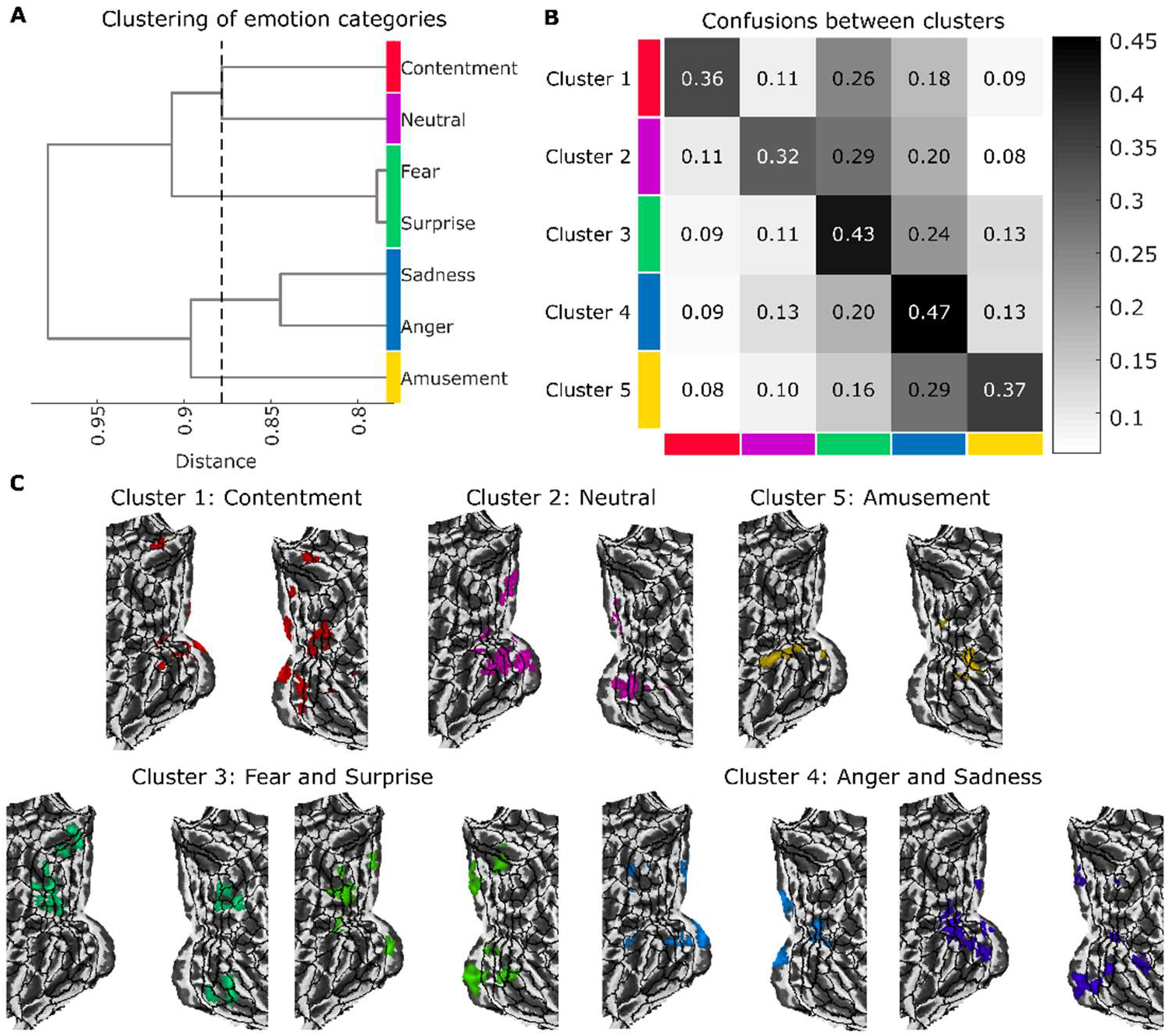
Multiclass classification of occipital lobe activity reveals five discriminable emotion clusters. (a) Dendrogram illustrates hierarchical clustering of emotion categories that maximizes discriminability. The x-axis indicates the inner squared distance between emotion categories. The dashed line shows the optimal clustering solution; cluster membership is indicated by color. (b) Confusion matrix for the five-cluster solution depicts the proportion of trials that are classified as belonging to each cluster (shown by the column) as a function of ground truth membership in a cluster (indicated by the row). The overall five-way accuracy is 40.54%, where chance is 20%. (c) Model weights indicate where increasing brain activity is associated with the prediction of each emotion category. Maps are thresholded at a voxel-wise threshold of *P* < .05 for display.

## Discussion

Our work demonstrates the intimate relationship between visual perception and emotion. Even though emotions are often about specific objects, events, or situations (Tooby and Cosmides), few accounts of emotion specify how sensory information is transformed into emotion-relevant signals in a computationally explicit fashion (Bach and Dayan 2017). Driven by the hypothesis that emotion schemas are embedded in the human visual system, we developed a computational model (EmoNet) to classify images into 20 different emotion categories. Consistent with our prediction that image features *alone* are sufficient for predicting normative ratings of emotion categories determined by humans, EmoNet accurately classified images into at least 11 different emotion categories in hold-out test data. Supporting our second prediction that EmoNet representations should map primarily onto the activity of sensory systems (as opposed to subcortical structures or limbic brain regions) distributed patterns of human occipital lobe activity were the best predictors of emotion category units in EmoNet. Finally, our third prediction was supported by the observation that patterns of occipital lobe activity were sufficient for decoding at least 15 emotion categories evoked by images, and at least five of seven emotional states elicited by cinematic movies. These findings both shed light on how visual processing constrains emotional responses, and how emotions are represented in the brain.

A large body of research has assumed that “low-level” visual information is mainly irrelevant to emotional processing; it should either be controlled for or explained away, even though studies have shown that neurons in early visual areas are sensitive to affective information such as reward (Shuler and Bear 2006). Our model provides a means to disentangle the visual properties of stimuli that are emotion-relevant from those that are not, and isolate stimulus-related features (e.g., red color serving as an indicator of higher energy content in fruit (Regan, Julliot et al. 2001, Melin, Chiou et al. 2017)) from more abstract constructs (e.g., the broader concept of ‘food craving’, which does not require a visual representation). Based on our findings, it seems unlikely that a complete account of emotion will be devoid of sensory qualities that are naturally associated with emotional outcomes, or those that are reliably learned through experience and influenced by culture.

We found that human ratings of pleasantness and excitement evoked by images can be accurately modeled as a combination of emotion-specific features (e.g., a mixture of features related to ‘disgust,’ ‘horror,’ ‘sadness,’ and ‘fear’ are highly predictive of unpleasant arousing experiences). Individuals may draw from this visual information when asked to rate images. The presence of emotion-specific visual features could activate learned associations with more general feelings of valence and arousal and help guide self-report. It is possible that feelings of valence and arousal arise from integration across feature detectors or predictive coding about the causes of interoceptive events, (Seth 2013, Barrett 2017). Rather than being irreducible (Barrett and Bliss-Moreau 2009), these feelings may be constructed from emotionally-relevant sensory information (Lindquist, Satpute et al. 2015), such as the emotion-specific features we have identified here, and prior expectations of their affective significance.

In addition to our observation that emotion-specific visual features can predict normative ratings of valence and arousal, we found that they were effective at classifying the genre of cinematic movie trailers. Moreover, the emotions that informed prediction were generally consistent with those typically associated with each genre (e.g., romantic comedies were predicted by activation of ‘romance’ and ‘amusement’). This validation differed from our other two image-based assessments of EmoNet (i.e., testing on hold-out videos from the database used for training, and testing on IAPS images) because it examined stimuli that are not conventionally used in the laboratory, yet are robust elicitors of emotional experience in daily life. Beyond hinting at real-world applications of our model, integrating results across these three validation tests serves to triangulate our findings (Lawlor, Tilling et al. 2016, Munafo and Davey Smith 2018), as different methods (with different assumptions and biases) were used to produce more robust, reproducible results.

The fact that emotion category units of EmoNet were best characterized by activity spanning visual cortex (i.e., the occipital lobe) sheds light on the nature of emotion representation in the brain, providing evidence for a distributed rather than a modular neural basis of emotion schemas. Activation of schemas in visual cortex offers a rapid, possibly automatic way of triggering downstream emotional responses in the absence of deliberative or top-down conceptual processes. By harnessing the parallel and distributed architecture of the visual system, these representations could be refined through experience. Information from downstream systems via feedback projections from ventromedial prefrontal cortex or the amygdala (Pessoa and Adolphs 2010, Kravitz, Saleem et al. 2013) could update visual emotion schemas through learning (Serences 2008, Dunsmoor, Kragel et al. 2014). Thus, emotion-related activity in visual cortex is most likely not a purely bottom-up response to stimuli or a top-down interpretation of them, but is at the interface of sensory representations of the environment and prior knowledge about potential outcomes. Future work integrating computational models with recurrent feedback (e.g., Nayebi, Bear et al. 2018) and brain responses to emotional images will be necessary to understand the convergence of bottom-up and top-down signals.

Our computational framework provides a way to resolve outstanding theoretical debates in affective science. It could be used, for example, to test if mappings between visual features and emotions are conserved across species or change throughout development in humans. Based on evolutionary accounts that suggest certain basic emotions are solutions to survival challenges, mechanisms for detecting emotionally significant events should be conserved across species. Notably, some of the most accurately predicted schemas include ‘sexual desire’ and ‘craving’ which are motivational states that transcend cultures and are linked to clear evolutionary goals (i.e., to reproduce and to acquire certain nutrients). Work in the domain of object recognition has shown that representations of objects are highly similar between humans and macaques (Kriegeskorte, Mur et al. 2008), an extension of the present work is to test whether the emotion representations we identified here are as well.

Our work has several limitations that can be addressed in future work. Although our goal was to focus on visual processing of emotional features, visual stimulation is not the only way in which emotions can be elicited. Information from other senses (olfactory, auditory, somatic, interoceptive, etc.), memories of past events, manipulation of motor activation, and mental imagery have all been used to evoke emotional experiences in the lab. EmoNet can be expanded, potentially by adding more abstract or ‘supramodal’ representation of emotions (Peelen, Atkinson et al. 2010, Skerry and Saxe 2014, Kim, Shinkareva et al. 2017) and interactions among different types of sensory information. Incorporating other, more neurally informed mechanisms into the model, such as recurrence and learning rules that are biologically plausible are possible directions for future development.

Using a combination of computational and neuroscientific tools, we have demonstrated that emotion schemas are embedded in the human visual system. By precisely specifying what makes images emotional, our modeling framework offers a new approach to understanding how visual inputs can rapidly evoke complex emotional responses. We anticipate that developing biologically inspired computational models will be a crucial next step for resolving debates about the nature of emotions (e.g., Adolphs 2017, Barrett 2017, Adolphs and Andler 2018) and providing practical tools for scientific research and in applied settings.

## Author contributions

conceptualization, P.A.K. and T.D.W.; investigation, P.A.K. and M.R.; methodology, P.A.K., M.R., K.S.L., and T.D.W.; software, P.A.K. and T.D.W.; writing – original draft, P.A.K., M.R., K.S.L., and T.D.W.; writing – review and editing P.A.K., M.R., K.S.L., and T.D.W.; visualization, P.A.K., supervision, T.D.W.

## Data and code availability

fMRI data are available at https://www.neurovault.org, analysis code is available at https://github.com/canlab

## Materials and Methods

### Computational model development

We used a large database of emotional video clips (Cowen and Keltner 2017) for developing EmoNet. This database includes 2,185 videos that are well characterized by 27 distinct emotion categories. A total of 137,482 frames were extracted from the videos and divided into training and testing samples using a 90-10 split. Emotion categories that had fewer than 1,000 frames for training were excluded from the model, reducing the emotions included in the model to ‘adoration’, ‘aesthetic appreciation’, ‘amusement’, ‘anxiety’, ‘awe’, ‘boredom’, ‘confusion’, ‘craving’, ‘disgust’, ‘empathic pain’, ‘entrancement’, ‘excitement’, ‘fear’, ‘horror’, ‘interest’, ‘joy’, ‘romance’, ‘sadness’, ‘sexual desire’, and ‘surprise’. The pre-trained CNN model AlexNet (Krizhevsky, Sutskever et al. 2012) was downloaded for use in MATLAB via MatConvNet (Vedaldi and Lenc 2015). We fixed all but the last fully-connected layer of AlexNet, and we retrained the model after replacing the 1,000 target object categories with the 20 emotion categories listed above. Training was performed using stochastic gradient descent with momentum, an initial learning rate of 0.0001, and a mini-batch size of 16.

### Computational model validation

Three separate tests were performed to assess model performance: 1) validation on the hold-out dataset, 2) predicting normative ratings of valence and arousal for the International Affective Picture System (IAPS, a standardized set of affective images used in psychological research (Lang, Bradley et al. 2008)), and 3) predicting the genre of cinematic movie trailers.

For the hold-out dataset, we computed standard signal detection metrics (i.e., AUC, sensitivity, and specificity) and evaluated overall model performance and that for each category. We performed inference on model performance by generating null distributions through random permutation of test-set labels. Additionally, EmoNet’s performance was compared to that of AlexNet to determine how much retraining the last fully-connected layer improved performance. For this purpose, we randomly sampled AlexNet predictions for 20 object categories to compute relevant signal detection metrics 10,000 times in addition to finding the 20 unique object categories that best predicted the 20 emotions.

We assessed the generalizability of EmoNet on IAPS images by using activations in the last fully-connected layer to predict normative ratings of valence and arousal. This analysis was performed using Partial Least Squares regression (with bootstrap procedures to estimate the variance of parameter estimates), and ten iterations of 10-fold cross-validation (Bouckaert and Frank 2004) to determine the correlation between model predictions and ‘ground truth’ normative ratings. We averaged normative ratings and EmoNet predictions for each of 25 quantiles. The construct validity of model parameters (e.g., whether greater activations of ‘amusement,’ as opposed to ‘fear,’ were associated with higher valence norms) and cross-validated estimates of root mean square error served as outcomes of interest.

In the final validation test, we used activations in the last fully-connected layer to classify the genre of movie trailers (*N* = 28, sampling from romantic comedy, horror, and action movies; see Appendix I). Trailers were selected based on genres listed on https://www.imdb.com/feature/genre/ and their availability at http://www.hd-trailers.net/. Classification into different genres was performed using Partial Least Squares regression (with bootstrap procedures to estimate the variance of parameter estimates), and 10-fold cross-validation to estimate the accuracy of classification into different genres. The construct validity of model parameters (e.g., whether greater activations of ‘amusement’ predicted romantic comedies) and cross-validated estimates of classification accuracy served as outcomes of interest.

### fMRI experiment I: modeling brain responses to emotional images

#### Participants

We recruited eighteen healthy, right-handed individuals (10 Female, *M*_age_ = 25) from the Boulder area. As there were, to our knowledge, no prior studies relating activation in convolutional neural nets to human fMRI responses to emotional images, this sample size was not determined a priori. The experimental design focused on maximizing task-related signal within subjects by showing participants 112 affective images. Confirmatory post-hoc analysis of effect size and the variance of parameter estimates corroborated that this sample size was sufficient for reliably detecting effects and minimizing the variance of parameter estimates (e.g., predicting EmoNet outcomes from occipital lobe activity using a random sample of only 9 participants produced an average effect size of *d* = 3.08, 95% CI = [2.08 4.36], see **Figure S4**). Participants did not meet DSM V criteria for any psychological disorder and were screened to ensure safety in the MR environment. All participants provided informed consent before the experiment in accordance with the University of Colorado Boulder Institutional Review Board

#### Experimental paradigm

In this experiment, brain activity was measured using fMRI while participants viewed a series of emotional images. Stimuli were selected from the IAPS and the Geneva Affective PicturE Database (GAPED) using published normative arousal ratings, to have either positive or negative valence and high arousal (Mikels, Fredrickson et al. 2005, Libkuman, Otani et al. 2007, Lang, Bradley et al. 2008, Dan-Glauser and Scherer 2011). A total of 112 images were used for this experiment.

Image presentation lasted 4s, with a jittered inter-trial-interval of 3 to 8-seconds (average ISI = 4s). The scanning session was divided into two runs lasting 7.5 minutes, where the images were presented in a randomized order. Stimulus presentation was controlled using code written in MATLAB using the Psychophysics toolbox extension (Brainard 1997, Kleiner, Brainard et al. 2007).

#### MRI data acquisition

Gradient-echo echo-planar imaging BOLD-fMRI was performed on a 3 Tesla Siemens MRI scanner (Siemens Healthcare). Functional images were acquired using Multiband EPI sequence: echo time = 30 ms, repetition time = 765 ms, flip angle = 44°, number of slices = 80, slice orientation = coronal, phase encoding = h > f, voxel size = 1.6 × 1.6 × 2.0 mm, gap between slices = 0 mm, field of view = 191 × 191 mm^2^, Multi-band acceleration factor = 8; echo spacing = 0.72 ms, bandwidth = 1,724 Hz per pixel, partial Fourier in the phase encode direction: 7/8.

Structural images were acquired using a single shot T1 MPRAGE sequence: echo time = 2.01 ms, repetition time = 2.4 s, flip angle = 8°, number of slices = 224, slice orientation = sagittal, voxel size = 0.8 mm isotropic, gap between slices = 0 mm, field of view = 256 × 256 mm^2^, GRAPPA acceleration factor = 2; echo spacing = 7.4 ms, bandwidth = 240 Hz per pixel.

#### MRI preprocessing

Multiband brain imaging data were preprocessed following procedures used in the Human Connectome Project (Glasser, Sotiropoulos et al. 2013). This approach includes distortion correction, spatial realignment based on translation (in the transverse, sagittal, and coronal planes) and rotation (roll, pitch, and yaw), spatial normalization to MNI152 space using T1 data, and smoothing using a 6mm FWHM Gaussian kernel. Preprocessing was completed using the Mind Research Network’s Auto-Analysis software (Bockholt, Scully et al. 2010).

#### MRI analysis

Preprocessed fMRI data were analyzed using general linear models with SPM 8 software (Wellcome Trust Centre for Neuroimaging, UK). Separate models were estimated for each participant that included: 1) a regressor for every image presented to subjects, modeled as a 4s boxcar convolved with the canonical hemodynamic response function of SPM, 2) 24 motion covariates from spatial realignment (i.e., translation in x, y, and z dimensions; roll, pitch, and yaw; and their first and second order temporal derivatives), 3) nuisance regressors specifying outlier timepoints, or ‘spikes’, that had large deviations in whole-brain BOLD signal, and 4) constant terms to model the mean of each imaging session.

To identify mappings between patterns of brain activity and features of EmoNet, partial least squares (PLS) regression models were fit on data from the entire sample (*N* = 18) using the full set of single-trial parameter estimates (112 trials for each subject) as input and activation in the last fully-connected layer of EmoNet as the output (20 different variables, one per emotion category). Model generalization (indicated by the correlation between observed and predicted outcomes and mean squared error) was estimated using leave-one-subject-out cross-validation. Inference on model performance was performed through permutation testing, where model features (i.e., activation in layer fc8) were randomly shuffled on each of 10,000 iterations. Performance relative to the noise ceiling was estimated by computing the ratio of cross-validated estimates to those using resubstitution (which should yield perfect performance in a noiseless setting; see Supplementary Text).

Inference on parameter estimates from PLS was performed via bootstrap resampling with 1,000 replicates, using the mean and standard error of the bootstrap distribution to compute *P*-values based on a normal distribution. Bootstrap distributions were visually inspected to verify that they were approximately normal. Thresholding of maps was performed using False Discovery Rate (FDR) correction with a threshold of *q* < .05. To visualize all 20 models in a low dimensional space, principal component decomposition was performed on PLS regression coefficients on every bootstrap iteration to produce a set of orthogonal components and associated coefficients comprising a unique pattern of occipital lobe voxels. Procedures for inference and thresholding were identical to those used for parameter estimates, only they were applied to coefficients from the PCA. Brain maps in the main figures are unthresholded for display. All results reported in the main text of the manuscript (and supplementary figures) survive FDR correction for multiple comparisons.

### fMRI experiment II: classifying brain responses to emotional film clips

fMRI data used for validating the model have been published previously; here we briefly summarize the procedure. Full details can be found in Kragel et al. (Kragel and LaBar 2015).

#### Participants

We used the full sample (*N* = 32) from an archival dataset characterizing brain responses to emotional films and music clips. For this analysis, which focuses on visual processing, we used only brain responses to film stimuli (available at http://www.neurovault.org). These data comprise single-trial estimates of brain activity for stimuli used to evoke experiences that were rated as being emotionally neutral in addition to states of contentment, amusement, surprise, fear, anger, and sadness.

#### Experimental paradigm

Participants completed an emotion induction task where they were presented with an emotional stimulus and subsequently provided on-line self-reports of emotional experience. Each trial started with the presentation of either a film or music clip (mean duration = 2.2 minutes), immediately followed by a 23-item affect self-report scale (Stephens, Christie et al. 2010) lasting 1.9 min followed by a 1.5 min washout clip to minimize carry-over effects.

#### MRI data acquisition

Scanning was performed on a 3 Tesla General Electric MR 750 system with 50-mT/m gradients and an eight-channel head coil for parallel imaging (General Electric, Waukesha, WI, USA). High-resolution images were acquired using a 3D fast SPGR BRAVO pulse sequence: repetition time (TR) = 7.58 ms; echo time (TE) = 2.936 ms; image matrix = 256^2^; α = 12°; voxel size = 1 × 1 × 1 mm; 206 contiguous slices. These structural images were aligned in the near-axial plane defined by the anterior and posterior commissures. Whole-brain functional images were acquired using a spiral-in pulse sequence with sensitivity encoding along the axial plane (TR = 2000 ms; TE = 30 ms; image matrix = 64 × 128, α = 70°; voxel size = 3.8 × 3.8 × 3.8 mm; 34 contiguous slices).

#### MRI preprocessing

fMRI data were preprocessed using SPM8 (http://www.fil.ion.ucl.ac.uk/spm). Images were first realigned to the first image of the series using a six-parameter, rigid-body transformation. The realigned images were then coregistered to each participant’s T1-weighted structural image and normalized to MNI152 space using high-dimensional warping implemented in the VBM8 toolbox. No additional smoothing was applied to the normalized images.

#### MRI analysis

A univariate general linear model (GLM) was used to create images for the prediction analysis. The model included separate boxcar regressors indicating the onset times for each stimulus, which allowed us to isolate responses to each emotion category. Separate regressors for the rating periods were included in the model but were not of interest. All regressors were convolved with the canonical HRF used in SPM, and an additional six covariate regressors modeled for movement effects.

Pattern classification of occipital lobe responses to the film clips was performed using Partial Least Squares Discriminant Analysis (PLS-DA; following methods in ref. (Kragel and LaBar 2015)). The data comprised 444 trials total (2 videos x 7 emotion categories x 32 subjects, with four trials excluded due to technical issues during scanning). Measures of classification performance were estimated using 8-fold subject independent cross-validation, where subjects were randomly divided into eight groups; classification models were iteratively trained on data from all but one group, and model performance was assessed on data from the hold-out group. This procedure was repeated until all data had been used for training and testing (8 folds total). Inference on model performance was made using permutation tests, where the above cross-validation procedure was repeated 1,000 times with randomly permuted class labels to produce a null distribution for inference. The number of emotion categories that could be accurately discriminated from one another was estimated using Discriminable Cluster Identification (see Supplementary Text for details). Inference on model weights (i.e., PLS parameter estimates) at each voxel was made via bootstrap resampling with a normal approximated interval.

#### Definition of regions-of-interest (ROIs)

A region-of-interest (ROI) approach was used to restrict features for model development and to localize where information about emotions is encoded. We selected several anatomically defined ROIs based on our focus on the visual system. These regions include multiple cytoarchitecturally defined visual areas (i.e., V1, V2, V3v, V3d, V3a, and V4 (Amunts, Malikovic et al. 2000, Rottschy, Eickhoff et al. 2007)), the entire occipital lobe (Lancaster, Woldorff et al. 2000), and inferotemporal cortex (Tzourio-Mazoyer, Landeau et al. 2002). These masks were created using the SPM Anatomy toolbox (Eickhoff, Stephan et al. 2005) and the Automated Anatomical Labeling atlas (Tzourio-Mazoyer, Landeau et al. 2002).

## References

Adolphs, R. (2013). “The biology of fear.” Curr Biol 23(2): R79–93.

Adolphs, R. (2017). “How should neuroscience study emotions? by distinguishing emotion states, concepts, and experiences.” Social Cognitive and Affective Neuroscience 12(1): 24–31.

Adolphs, R. and D. Andler (2018). “Investigating Emotions as Functional States Distinct From Feelings.” Emotion Review 10(3): 191–201.

Amunts, K., A. Malikovic, H. Mohlberg, T. Schormann and K. Zilles (2000). “Brodmann’s Areas 17 and 18 Brought into Stereotaxic Space—Where and How Variable?” NeuroImage 11(1): 66–84.

Bach, D. R. and P. Dayan (2017). “Algorithms for survival: a comparative perspective on emotions.” Nat Rev Neurosci 18(5): 311–319.

Barrett, L. F. (2006). “Solving the emotion paradox: categorization and the experience of emotion.” Pers Soc Psychol Rev 10(1): 20–46.

Barrett, L. F. (2017). “The theory of constructed emotion: an active inference account of interoception and categorization.” Soc Cogn Affect Neurosci 12(11): 1833.

Barrett, L. F. (2017). “The theory of constructed emotion: an active inference account of interoception and categorization.” Social Cognitive and Affective Neuroscience 12(1): 1–23.

Barrett, L. F. and M. Bar (2009). “See it with feeling: affective predictions during object perception.” Philosophical Transactions of the Royal Society of London B: Biological Sciences 364(1521): 1325–1334.

Barrett, L. F. and E. Bliss-Moreau (2009). “Affect as a psychological primitive.” Advances in experimental social psychology 41: 167–218.

Berridge, K. and P. Winkielman (2003). “What is an unconscious emotion?(The case for unconscious “liking”).” Cognition and emotion 17(2): 181–211.

Bockholt, H., M. Scully, W. Courtney, S. Rachakonda, A. Scott, A. Caprihan, J. Fries, R. Kalyanam, J. Segall, R. De La Garza, S. Lane and V. Calhoun (2010). “Mining the mind research network: a novel framework for exploring large scale, heterogeneous translational neuroscience research data sources.” Frontiers in Neuroinformatics 3(36).

Bouckaert, R. R. and E. Frank (2004). Evaluating the Replicability of Significance Tests for Comparing Learning Algorithms, Berlin, Heidelberg, Springer Berlin Heidelberg.

Bower, G. H. (1981). “Mood and memory.” American Psychologist 36(2): 129–148.

Bradley, M. M. and P. J. Lang (1999). Affective norms for English words (ANEW): Instruction manual and affective ratings, Citeseer.

Brainard, D. H. (1997). “The psychophysics toolbox.” Spatial vision 10: 433–436.

Carroll, J. B. (1953). “An analytical solution for approximating simple structure in factor analysis.” Psychometrika 18(1): 23–38.

Chang, L. J., P. J. Gianaros, S. B. Manuck, A. Krishnan and T. D. Wager (2015). “A Sensitive and Specific Neural Signature for Picture-Induced Negative Affect.” PLoS Biol 13(6): e1002180.

Cowen, A. S. and D. Keltner (2017). “Self-report captures 27 distinct categories of emotion bridged by continuous gradients.” Proceedings of the National Academy of Sciences of the United States of America 114(38): E7900–E7909.

Dahnke, R., R. A. Yotter and C. Gaser (2013). “Cortical thickness and central surface estimation.” Neuroimage 65: 336–348.

Dan-Glauser, E. S. and K. R. Scherer (2011). “The Geneva affective picture database (GAPED): a new 730-picture database focusing on valence and normative significance.” Behavior Research Methods 43(2): 468.

Dunsmoor, J. E., P. A. Kragel, A. Martin and K. S. LaBar (2014). “Aversive learning modulates cortical representations of object categories.” Cereb Cortex 24(11): 2859–2872.

Eickhoff, S. B., K. E. Stephan, H. Mohlberg, C. Grefkes, G. R. Fink, K. Amunts and K. Zilles (2005). “A new SPM toolbox for combining probabilistic cytoarchitectonic maps and functional imaging data.” NeuroImage 25(4): 1325–1335.

Ekman, P. (1992). “An Argument for Basic Emotions.” Cognition & Emotion 6(3-4): 169–200.

Ekman, P. and D. Cordaro (2011). “What is Meant by Calling Emotions Basic.” Emotion Review 3(4): 364–370.

Eradath, M. K., T. Mogami, G. Wang and K. Tanaka (2015). “Time Context of Cue-Outcome Associations Represented by Neurons in Perirhinal Cortex.” The Journal of Neuroscience 35(10): 4350–4365.

Esterman, M., B. J. Tamber-Rosenau, Y.-C. Chiu and S. Yantis (2010). “Avoiding non-independence in fMRI data analysis: Leave one subject out.” NeuroImage 50(2): 572–576.

Fontaine, J. R., K. R. Scherer, E. B. Roesch and P. C. Ellsworth (2007). “The world of emotions is not two-dimensional.” Psychological science 18(12): 1050–1057.

Gaser, C. and R. Dahnke (2016). “CAT-a computational anatomy toolbox for the analysis of structural MRI data.” HBM 2016: 336–348.

Glasser, M. F., T. S. Coalson, E. C. Robinson, C. D. Hacker, J. Harwell, E. Yacoub, K. Ugurbil, J. Andersson, C.F. Beckmann, M. Jenkinson, S. M. Smith and D. C. Van Essen (2016). “A multi-modal parcellation of human cerebral cortex.” Nature 536: 171.

Glasser, M. F., S. N. Sotiropoulos, J. A. Wilson, T. S. Coalson, B. Fischl, J. L. Andersson, J. Xu, S. Jbabdi, M. Webster, J. R. Polimeni, D. C. Van Essen and M. Jenkinson (2013). “The minimal preprocessing pipelines for the Human Connectome Project.” NeuroImage 80: 105–124.

Haenny, P. and P. Schiller (1988). “State dependent activity in monkey visual cortex.” Experimental Brain Research 69(2): 225–244.

Held, R. and A. Hein (1963). “Movement-produced stimulation in the development of visually guided behavior.” Journal of comparative and physiological psychology 56(5): 872.

Izard, C. E. (2007). “Basic Emotions, Natural Kinds, Emotion Schemas, and a New Paradigm.” Perspectives on Psychological Science 2(3): 260–280.

Kahneman, D. and P. Egan (2011). Thinking, fast and slow, Farrar, Straus and Giroux New York.

Kim, J., S. V. Shinkareva and D. H. Wedell (2017). “Representations of modality-general valence for videos and music derived from fMRI data.” NeuroImage 148: 42–54.

Kleiner, M., D. Brainard, D. Pelli, A. Ingling, R. Murray and C. Broussard (2007). “What’s new in Psychtoolbox-3.” Perceptions.

Kohavi, R. (1995). A study of cross-validation and bootstrap for accuracy estimation and model selection. Ijcai, Stanford, CA.

Kragel, P. A., L. Koban, L. F. Barrett and T. D. Wager (2018). “Representation, Pattern Information, and Brain Signatures: From Neurons to Neuroimaging.” Neuron 99(2): 257–273.

Kragel, P. A. and K. S. LaBar (2015). “Multivariate neural biomarkers of emotional states are categorically distinct.” Soc Cogn Affect Neurosci 10(11): 1437–1448.

Kravitz, D. J., K. S. Saleem, C. I. Baker, L. G. Ungerleider and M. Mishkin (2013). “The ventral visual pathway: An expanded neural framework for the processing of object quality.” Trends in cognitive sciences 17(1): 26–49.

Kriegeskorte, N., M. Mur, D. A. Ruff, R. Kiani, J. Bodurka, H. Esteky, K. Tanaka and P. A. Bandettini (2008). “Matching categorical object representations in inferior temporal cortex of man and monkey.” Neuron 60(6): 1126–1141.

Krishnan, A., C. W. Woo, L. J. Chang, L. Ruzic, X. Gu, M. Lopez-Sola, P. L. Jackson, J. Pujol, J. Fan and T. D. Wager (2016). “Somatic and vicarious pain are represented by dissociable multivariate brain patterns.” Elife 5.

Krizhevsky, A., I. Sutskever and G. E. Hinton (2012). Imagenet classification with deep convolutional neural networks. Advances in neural information processing systems.

Kurdi, B., S. Lozano and M. R. Banaji (2017). “Introducing the Open Affective Standardized Image Set (OASIS).” Behavior Research Methods 49(2): 457–470.

Lancaster, J. L., M. G. Woldorff, L. M. Parsons, M. Liotti, C. S. Freitas, L. Rainey, P. V. Kochunov, D. Nickerson, S. A. Mikiten and P. T. Fox (2000). “Automated Talairach atlas labels for functional brain mapping.” Hum Brain Mapp 10(3): 120–131.

Lang, P. and M. M. Bradley (2007). “The International Affective Picture System (IAPS) in the study of emotion and attention.” Handbook of emotion elicitation and assessment 29.

Lang, P. J., M. M. Bradley and B. N. Cuthbert (2008). International affective picture system (IAPS): Affective ratings of pictures and instruction manual, University of Florida, Gainesville, FL.

Lawlor, D. A., K. Tilling and G. Davey Smith (2016). “Triangulation in aetiological epidemiology.” International Journal of Epidemiology 45(6): 1866–1886.

Lazarus, R. S. (1966). “Psychological stress and the coping process.”

Lazarus, R. S. (1968). Emotions and adaptation: Conceptual and empirical relations. Nebraska symposium on motivation, University of Nebraska Press.

Libkuman, T. M., H. Otani, R. Kern, S. G. Viger and N. Novak (2007). “Multidimensional normative ratings for the International Affective Picture System.” Behavior Research Methods 39(2): 326–334.

Lindquist, K. A., A. B. Satpute, T. D. Wager, J. Weber and L. F. Barrett (2015). “The brain basis of positive and negative affect: evidence from a meta-analysis of the human neuroimaging literature.” Cerebral Cortex 26(5): 1910–1922.

MacLean, P. D. (1952). “Some psychiatric implications of physiological studies on frontotemporal portion of limbic system (visceral brain).” Clinical Neurophysiology 4(4): 407–418.

Mayberg, H. S., M. Liotti, S. K. Brannan, S. McGinnis, R. K. Mahurin, P. A. Jerabek, J. A. Silva, J. L. Tekell, C. C. Martin, J. L. Lancaster and P. T. Fox (1999). “Reciprocal limbic-cortical function and negative mood: converging PET findings in depression and normal sadness.” Am J Psychiatry 156(5): 675–682.

McAlonan, K., J. Cavanaugh and R. H. Wurtz (2008). “Guarding the gateway to cortex with attention in visual thalamus.” Nature 456(7220): 391.

Melin, A. D., K. L. Chiou, E. R. Walco, M. L. Bergstrom, S. Kawamura and L. M. Fedigan (2017). “Trichromacy increases fruit intake rates of wild capuchins (Cebus capucinus imitator).” Proceedings of the National Academy of Sciences 114(39): 10402–10407.

Mikels, J. A., B. L. Fredrickson, G. R. Larkin, C. M. Lindberg, S. J. Maglio and P. A. Reuter-Lorenz (2005). “Emotional category data on images from the International Affective Picture System.” Behavior research methods 37(4): 626–630.

Mogami, T. and K. Tanaka (2006). “Reward Association Affects Neuronal Responses to Visual Stimuli in Macaque TE and Perirhinal Cortices.” The Journal of Neuroscience 26(25): 6761–6770.

Moors, A. (2018). Appraisal Theory of Emotion. Encyclopedia of Personality and Individual Differences. V. Zeigler-Hill and T. K. Shackelford. New York, Springer.

Morris, J. S., K. J. Friston, C. Büchel, C. D. Frith, A. W. Young, A. J. Calder and R. J. Dolan (1998). “A neuromodulatory role for the human amygdala in processing emotional facial expressions.” Brain 121(1): 47–57.

Munafo, M. R. and G. Davey Smith (2018). “Robust research needs many lines of evidence.” Nature 553(7689): 399–401.

Nayebi, A., D. Bear, J. Kubilius, K. Kar, S. Ganguli, D. Sussillo, J. J. DiCarlo and D. L. Yamins (2018). “Task-Driven Convolutional Recurrent Models of the Visual System.” arXiv preprint arXiv:1807.00053.

Niedenthal, P. M. (2007). “Embodying emotion.” Science 316(5827): 1002–1005.

O’Connor, D. H., M. M. Fukui, M. A. Pinsk and S. Kastner (2002). “Attention modulates responses in the human lateral geniculate nucleus.” Nature neuroscience 5(11): 1203.

Öhman, A. and S. Mineka (2003). “The malicious serpent: Snakes as a prototypical stimulus for an evolved module of fear.” Current directions in psychological science 12(1): 5–9.

Panksepp, J. (1998). Affective neuroscience: the foundations of human and animal emotions. New York, Oxford University Press.

Peelen, M. V., A. P. Atkinson and P. Vuilleumier (2010). “Supramodal representations of perceived emotions in the human brain.” J Neurosci 30(30): 10127–10134.

Pessiglione, M., B. Seymour, G. Flandin, R. J. Dolan and C. D. Frith (2006). “Dopamine-dependent prediction errors underpin reward-seeking behaviour in humans.” Nature 442(7106): 1042–1045.

Pessoa, L. (2008). “On the relationship between emotion and cognition.” Nature Reviews Neuroscience 9(2): 148–158.

Pessoa, L. and R. Adolphs (2010). “Emotion processing and the amygdala: from a ‘low road’ to ‘many roads’ of evaluating biological significance.” Nat Rev Neurosci 11(11): 773–783.

Plutchik, R. (1997). The circumplex as a general model of the structure of emotions and personality, American Psychological Association.

Rasheed, Z. and M. Shah (2002). Movie genre classification by exploiting audio-visual features of previews. Object recognition supported by user interaction for service robots.

Recanzone, G., C. Schreiner and M. Merzenich (1993). “Plasticity in the frequency representation of primary auditory cortex following discrimination training in adult owl monkeys.” The Journal of Neuroscience 13(1): 87–103.

Regan, B. C., C. Julliot, B. Simmen, F. Viénot, P. Charles–Dominique and J. D. Mollon (2001). “Fruits, foliage and the evolution of primate colour vision.” Philosophical Transactions of the Royal Society of London. Series B: Biological Sciences 356(1407): 229–283.

Rifkin, R. and A. Klautau (2004). “In Defense of One-Vs-All Classification.” J. Mach. Learn. Res. 5: 101–141.

Rottschy, C., S. B. Eickhoff, A. Schleicher, H. Mohlberg, M. Kujovic, K. Zilles and K. Amunts (2007). “Ventral visual cortex in humans: Cytoarchitectonic mapping of two extrastriate areas.” Human Brain Mapping 28(10): 1045–1059.

Russell, J. A. (1980). “A circumplex model of affect.” Journal of personality and social psychology 39(6): 1161.

Russell, J. A. (2003). “Core affect and the psychological construction of emotion.” Psychol Rev 110(1): 145–172.

Russell, J. A. and L. F. Barrett (1999). “Core affect, prototypical emotional episodes, and other things called emotion: dissecting the elephant.” Journal of personality and social psychology 76(5): 805.

Saarimaki, H., L. F. Ejtehadian, E. Glerean, I. P. Jaaskelainen, P. Vuilleumier, M. Sams and L. Nummenmaa (2018). “Distributed affective space represents multiple emotion categories across the human brain.” Soc Cogn Affect Neurosci 13(5): 471–482.

Saarimaki, H., A. Gotsopoulos, I. P. Jaaskelainen, J. Lampinen, P. Vuilleumier, R. Hari, M. Sams and L. Nummenmaa (2016). “Discrete Neural Signatures of Basic Emotions.” Cereb Cortex 26(6): 2563–2573.

Sasikumar, D., E. Emeric, V. Stuphorn and C. E. Connor (2018). “First-Pass Processing of Value Cues in the Ventral Visual Pathway.” Current Biology 28(4): 538–548.e533.

Scherer, K. R. (1984). On the nature and function of emotion: A component process approach. Approaches to emotion. K. R. Scherer and P. Ekman. Hillsdale, NJ: Erlbaum: 293–317.

Serences, J. T. (2008). “Value-based modulations in human visual cortex.” Neuron 60(6): 1169–1181.

Seth, A. K. (2013). “Interoceptive inference, emotion, and the embodied self.” Trends Cogn Sci 17(11): 565–573.

Shuler, M. G. and M. F. Bear (2006). “Reward Timing in the Primary Visual Cortex.” Science 311(5767): 1606–1609.

Skerry, A. E. and R. Saxe (2014). “A common neural code for perceived and inferred emotion.” J Neurosci 34(48): 15997–16008.

Stephens, C. L., I. C. Christie and B. H. Friedman (2010). “Autonomic specificity of basic emotions: Evidence from pattern classification and cluster analysis.” Biological Psychology 84(3): 463–473.

Tellegen, A., D. Watson and L. A. Clark (1999). “On the dimensional and hierarchical structure of affect.” Psychological Science 10(4): 297–303.

Tooby, J. and L. Cosmides (2008). “The evolutionary psychology of the emotions and their relationship to internal regulatory variables.”

Tzourio-Mazoyer, N., B. Landeau, D. Papathanassiou, F. Crivello, O. Etard, N. Delcroix, B. Mazoyer and M. Joliot (2002). “Automated Anatomical Labeling of Activations in SPM Using a Macroscopic Anatomical Parcellation of the MNI MRI Single-Subject Brain.” NeuroImage 15(1): 273–289.

Vedaldi, A. and K. Lenc (2015). Matconvnet: Convolutional neural networks for matlab. Proceedings of the 23rd ACM international conference on Multimedia, ACM.

Vuilleumier, P., M. P. Richardson, J. L. Armony, J. Driver and R. J. Dolan (2004). “Distant influences of amygdala lesion on visual cortical activation during emotional face processing.” Nature Neuroscience 7: 1271.

Warriner, A. B., V. Kuperman and M. Brysbaert (2013). “Norms of valence, arousal, and dominance for 13,915 English lemmas.” Behavior research methods 45(4): 1191–1207.

Yamins, D. L., H. Hong, C. F. Cadieu, E. A. Solomon, D. Seibert and J. J. DiCarlo (2014). “Performance-optimized hierarchical models predict neural responses in higher visual cortex.” Proc Natl Acad Sci U S A 111(23): 8619–8624.

Yamins, D. L. K. and J. J. DiCarlo (2016). “Using goal-driven deep learning models to understand sensory cortex.” Nature Neuroscience 19: 356.

Zajonc, R. B. (1984). “On the primacy of affect.”

